# Metacognitive Judgments during Visuomotor Learning Reflect the Integration of Error History

**DOI:** 10.1101/2023.01.17.524436

**Authors:** Christopher L. Hewitson, Naser Al-Fawakhiri, Alexander D. Forrence, Samuel D. McDougle

## Abstract

People form metacognitive representations of their own abilities across a range of tasks. How these representations are influenced by errors during learning is poorly understood. Here we ask how metacognitive confidence judgments of performance during motor learning are shaped by the learner’s recent history of errors. Across four motor learning experiments, our computational modeling approach demonstrated that people’s confidence judgments are best explained by a recency-weighted averaging of visually observed errors. Moreover, in the formation of these confidence estimates, people appear to re-weight observed motor errors according to a subjective cost function. Confidence judgments were adaptive, incorporating recent motor errors in a manner that was sensitive to the volatility of the learning environment, integrating a shallower history when the environment was more volatile. Finally, confidence tracked motor errors in the context of both implicit and explicit motor learning, but only showed evidence of influencing behavior in the latter. Our study thus provides a novel descriptive model that successfully approximates the dynamics of metacognitive judgments during motor learning.

**NEW & NOTEWORTHY:** This study examined how, during visuomotor-learning, people’s confidence in their performance is shaped by their recent history of errors. Using computational modeling, we found that confidence incorporated recent error-history, tracked subjective error-costs, was sensitive to environmental volatility, and in some contexts may influence learning. Together, these results provide a novel model of metacognitive judgments during motor-learning that could be applied to future computational and neural studies at the interface of higher-order cognition and motor control.

## Introduction

Humans have the ability to monitor qualities of their own performance in a task, a capacity often referred to as “metacognition.” Metacognitive processes have been observed across a range of domains, including simple perceptual decision-making (1), value-based decision-making (2), social cognition (3), and memory (4). Over a century of research has shown that people’s metacognitive judgements (e.g., their explicitly reported confidence in their choices/abilities) often closely track behavioral metrics like accuracy and response time (5). In one of the most studied laboratory models of metacognition and confidence – perceptual decision-making – researchers have used computational models to uncover strong links between confidence in a choice and the perceptual evidence accumulated for that choice over its competitors (e.g., ‘those dots are mostly moving left’) (6–8). Moreover, researchers have even discovered certain neural populations that simultaneously encode both accumulated evidence and decision confidence (9).

While studying confidence for independent choices (actions) has been a fruitful endeavor, one putatively more ecologically relevant question is how confidence evolves during the *learning* of actions (e.g., motor learning, reinforcement learning, etc.). Consider the experience of learning a new tune on an instrument: as one reduces their error, their mental representation of their own skill – a metacognitive judgment – evolves over time (e.g., “oh, I’m getting good at that section”). But instead of thinking of each action as an independent decision, as in perceptual decision-making tasks, one’s extended error history is likely key to determining how confident they are in specific actions.

While there has been minimal research on confidence and motor learning, there has been recent research on confidence and learning in nonmotor domains (3, 10–15). One recent study (10) used a perceptual decision task in which participants reported their estimate of the transition probabilities between two visual or auditory stimuli as well as their confidence in this report. The results indicated that participants not only learn a statistical model of transition probabilities over time, but also that their confidence ratings closely track this statistical inference. This work demonstrates that in a perceptual decision-making context, people’s confidence judgments closely correlate with their performance in tracking stochastic variables (16). Other work in the reinforcement learning domain has suggested that confidence in one’s choices during learning evolves along with learned latent value representations, and is subject to value-driven biases (2). Moreover, volatility in an environment, a second order statistic tracked over many trials, induces uncertainty in an agent, and agents tend to operate with a faster learning rate in these uncertain environments (17). This work suggests that higher-order variables like confidence may also correspond to the statistical uncertainty that underpins the learning process itself (18, 19).

Subjective confidence in the domain of motor learning has been less well studied, but some work has attempted to capture the role of continuous motor errors on subjective evaluations of confidence. For instance, one recent study (20) showed that individuals are able to predict their future performance on a sensorimotor task, and also leverage their confidence in their future performance to maximize future rewards. Another recent study (21) demonstrated that subjective confidence tracks one’s precision in a continuous temporal estimation task. Some work on motor sequence learning has looked at a more ‘zoomed-out’ form of confidence – block- and day-level judgments of one’s own ability (22). Lastly, relevant recent computational work has shown that individuals might utilize information about their prior motor variability to make confidence judgments of their motor precision, both prospectively (before movement, as we do in this study) and retrospectively (after movement) (23).

While these studies suggest that confidence in a motor context integrates prior history of performance (perhaps in a Bayesian manner), they do not directly address metacognitive dynamics during the protracted alteration of motor commands (24), the context of interest here. We believe studying confidence during motor learning may be key to understanding how people actively monitor their own performance, how the brain might represent uncertainty in movements over time, and perhaps how confidence may modulate the gain on the motor system’s response to errors. We thus address the question of how confidence evolves during motor learning, which may shed light on the psychological processes involved in motor learning, computationally isolate higher-level metacognitive variables for investigation in future neural studies, and perhaps be useful for increasing people’s motivation to learn in clinical and non clinical settings.

We performed four experiments and used a descriptive computational modeling approach to study confidence during motor learning. We hypothesized that heuristic models that take into account one’s recent error history would sufficiently capture the evolution of confidence judgments during motor learning, and would do so in a manner that adapts to the distribution of errors. We compared these models to alternative hypotheses that modeled confidence as a simple “read-out” of the most recent error outcome. Given recent distinctions between cognitive versus “low-level” forms of motor learning (10), we also tested our models in cases where motor learning was more “explicit” (i.e., requiring deliberate changes in action selection; Experiments 1-2) versus implicit (i.e., involving sub-conscious re-calibration of movement kinematics; Experiments 3-4), allowing us to test the effects of different forms of motor learning on confidence. Finally, we also performed exploratory analyses asking if confidence in turn exerts an effect on (explicit or implicit) motor learning.

To preview our key findings, this straightforward model significantly outperforms other model variants that do not incorporate error history, and also reveals that the dynamics of confidence ratings during motor learning adapt to environmental volatility. The model succeeds in both explicit and implicit motor learning contexts, supporting the idea that confidence tracks a generic error signal (however, the specific dynamics of confidence differed between these two contexts). Finally, in an exploratory analysis we present evidence that confidence can also affect learning – in cases where learning was primarily driven by changes in explicit action selection, those changes were better explained when considering the effects of both motor error and confidence judgments versus just motor error alone.

## METHODS

### Participants

Experiments 1 and 2 were conducted in-lab. A total of 38 neurologically healthy participants (Experiment 1: N = 18; Age = 21±5 years; Gender: 10 identified as Female, 1 preferred not to answer; Handedness: 16 right-handed [>40 on the Edinburgh handedness inventory (25). Experiment 2: N = 20; Age = 20±5 years; Gender: 13 identified as Female; Handedness: 19 right-handed) from the Yale University subject pool participated in this study.

For Experiments 3 and 4, participants were recruited via Prolific, an online platform. Recruitment was restricted to residents of the United States who were fluent in English, had normal or corrected to normal vision, had a Prolific approval rating greater than 95%, and had completed at least 40 web-based experiments previously. Legal sex (the current designation on a person’s birth certificate, driver’s license, and/or U.S. state identification) but not gender-identity was reported on this platform. Handedness was self-reported within the categories of right-handed, left-handed, or ambidextrous. A total of 55 participants (Experiment 3: N = 25; Age = 28±5 years; Sex: 10 Females; Handedness: 24 right-handed, 1 ambidextrous.

Experiment 4: N = 30 Age = 27±5 years; Sex: 13 Female; Handedness: 28 right-handed, 2 ambidextrous) performed the online experiments. In-lab participants received monetary compensation or course credit for their participation. Online participants received monetary compensation. Written informed consent was obtained from all participants before testing and the experimental protocol was approved in accordance with Yale University’s Institutional Review Board. Online participants who did not vary their confidence ratings over the course of more than 20 consecutive trials were excluded. This excluded 1 participant from Experiment 3 and 2 participants from Experiment 4.

### Apparatus

For Experiments 1 and 2, participants sat on a height-adjustable chair facing a 24.5-in. LCD monitor (Asus VG259QM; display size: 543.74 mm x 302.62 mm; resolution: 1920 x 1080 pixels; frame rate set to 240Hz; 1 ms response time), positioned horizontally ∼30 cm in front of the participant above the table platform, thus preventing direct vision of the hand (Fig. 1A). In their dominant hand they held a stylus embedded within a custom modified paddle, which they could slide across a digitizing tablet (Wacom PTH860; active area: 311 mm x 216 mm). Hand position was recorded from the tip of the stylus sampled by the tablet at 200 Hz. Stimulus presentation and movement recording were controlled by a custom-built Octave script (GNU Octave v5.2.0; Psychtoolbox-3 v3.0.18; Ubuntu 20.04.4 LTS). Aiming and confidence ratings were controlled by the non-dominant hand and entered on a USB keyboard (Fig. 1B).

**Figure 1.**
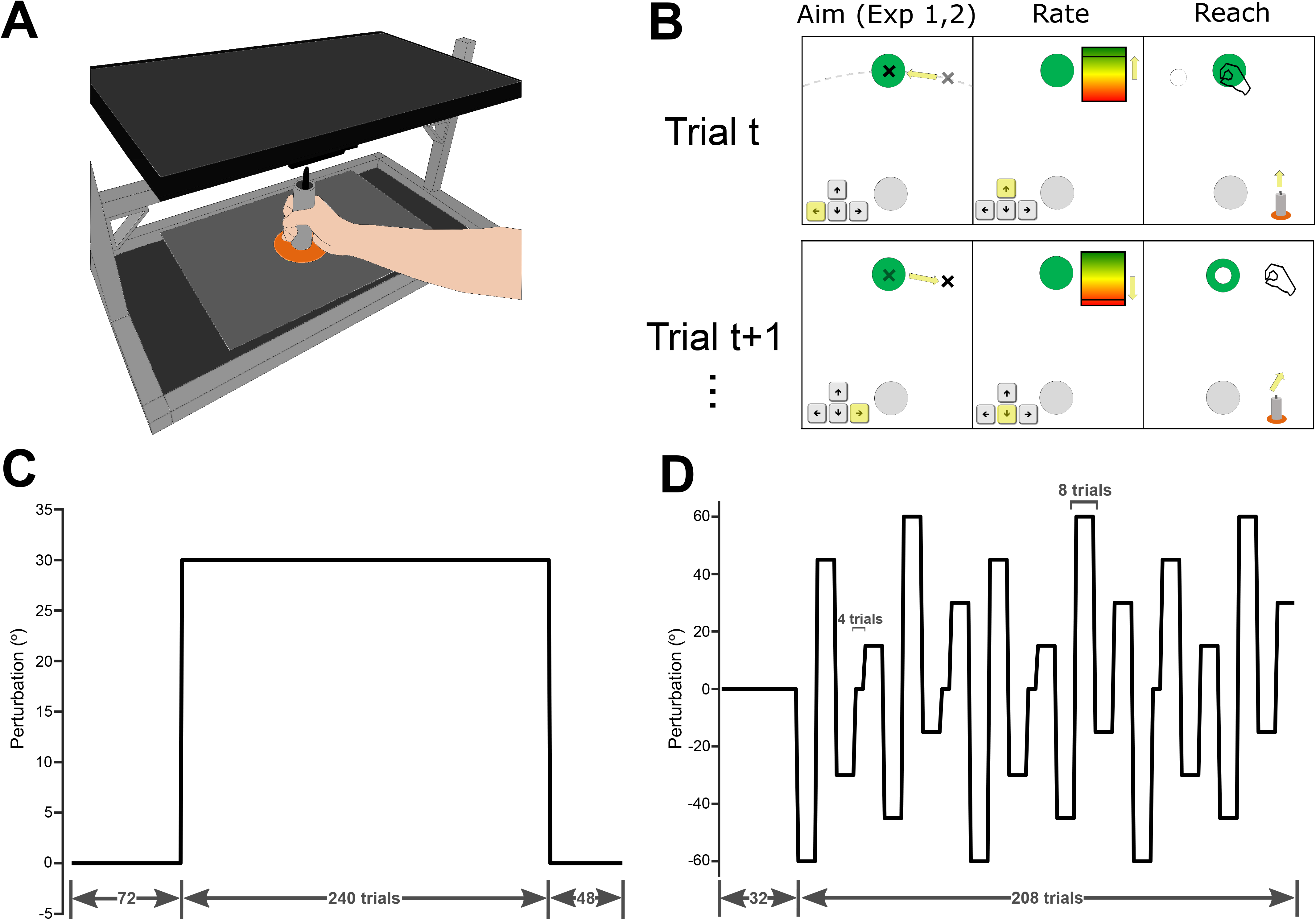
Experimental Design. (A) Experimental apparatus. Participants made reaching movements over a digitizing tablet while holding a modified air hockey paddle. (B) Schematic of two example trials. In Experiments 1 and 2 only, participants first moved a crosshair to specify their intended reach direction (“Aim”), then in all experiments, participants rated their confidence in that their movement would succeed in striking the target (“Report”), and finally executed their reach with feedback (“Reach”). (C) Perturbation schedule in Experiment 1. (D) Perturbation schedule in Experiment 2 (E) Perturbation schedule in Experiment 3. (D). Perturbation schedule in Experiment 4. The gray bars in D and E represent the range of perturbation values pseudo-randomly drawn on each trial (mean ±7.5°).

For Experiments 3 and 4, participants were restricted to use their home desktop computers with a mouse and keyboard. Average monitor frame-rates were 90Hz (SD: 40Hz) [60Hz (54%),120Hz (20%), 165Hz (12%), 144Hz (10%), 75Hz (4%)]. Stimulus presentation and movement recording were controlled by custom-built JavaScript code (23). The online task environment differed in terms of the inclusion of on-screen instructions at the start of the task, in place of verbal instructions provided in-lab.

### Task

Trials in Experiments 1 and 2 consisted of an aiming phase, a confidence rating phase, and then a reaching phase (Fig. 1B). The aiming phase was removed in Experiments 3 and 4. To briefly summarize the phases: During the aiming phase, participants were instructed to “position the aiming reticle where you intend to move your hand”; during the confidence reporting phase, participants were instructed to “rate how confident you are that where you aimed is correct” (i.e., that you would succeed at hitting the target given that movement plan); and during the reach phase, subjects made rapid reaches to the displayed target.

During the reaching phase, participants performed center-out-reaching movements from a central start location in the center of the monitor to try and accurately slice through a visual cursor on one of 8 visual targets (0°, 45°, 90°, 135°, 180°, 225°, 270°, and 315°) arranged around an invisible circle with a radius of 10 cm. In Experiments 1 and 2, the target location for each trial was pseudo-randomly selected. In Experiments 3 and 4, the target location was fixed at 45° for each trial. An off-axis target location was selected to avoid any perceptual and/or motor bias that may be present during reaches in a cardinal direction. Participants were instructed to move the stylus as quickly as possible from the start location in the direction of the displayed target (or the direction of their reported aiming location) and “slice through it.” The start location was marked by a filled white circle of 7 mm in diameter. The target locations were marked by filled green circles of 10 mm in diameter. Online visual feedback was given by a cursor (filled white circle, radius 2.5 mm), except for the washout phase in Experiment 1 (see below). If the reach duration exceeded 400 ms, a text prompt appeared on the monitor reminding participants to “please speed up your reach,” and the trial was repeated (with a new target location in Experiments 1 and 2).

During the aiming phase (Experiments 1 and 2 only), a white crosshair 7 mm in diameter was overlaid on the target (Figure 1B). Its movement was constrained to the invisible circle with a radius of 10 cm from the start location. The aiming crosshair’s location was adjusted with the left hand using the left and right arrow keys, which drove crosshair movements to the counter-clockwise and clockwise directions, respectively. When participants were satisfied with the match between their intended reach direction and the aiming crosshair’s position they registered their aim with the ‘enter’ key.

During the confidence rating phase, which directly followed the aiming phase when included, a rating bar (20mm x 40mm) was displayed 15° counterclockwise of the target. A white line, representing the participant’s confidence rating was initialized in the middle of the bar (50% confidence). As confidence increased towards 100%, the bar’s color changed from yellow to green. As confidence decreased towards 0%, the bar’s color changed from yellow to red. Participants reported their confidence level with their left hand using the up and down arrows, and registered their confidence rating with the ‘enter’ key.

Experiment 1 included reach baseline, report practice, adaptation, and washout blocks (Figure 1C). Baseline consisted of 24 trials (3 trials per target) with veridical online cursor feedback provided for the duration of the reach. Report practice consisted of 48 trials (6 trials per target) partitioned into the ‘aim’, confidence report,’ and ‘reach’ phases. Online visual feedback was provided throughout all reaches, except during the washout phase in Experiment 1 which had no visual feedback. Adaptation consisted of 240 trials (30 trials per target) that included all three trial phases (Figure 1C). Crucially, during the reach phase the cursor was rotated by 30° (with CW/CCW rotations evenly counterbalanced across participants). Washout (Experiment 1 only) consisted of 48 reach trials (6 trials per target) with no cursor feedback provided for the duration of the reach.

Experiment 2 included 16 baseline trials (2 per target), 16 report practice trials (2 per target), 208 adaptation trials (26 per target) and no washout trials (Figure 1D). The adaptation trials differed from Experiment 1 only in terms of the rotation perturbation applied to the cursor. In Experiment 2, rotation angles of -60°, -45°, -30°, -15°, 15°, 30°, 45°, and 60° were pseudo-randomly applied across 24 blocks of 8 trial mini-blocks (3 x 8 trial mini-blocks per rotation angle, across 8 targets, thus 192 rotation trials total). Four additional mini-blocks consisting of 4 trials of 0° rotation were interleaved throughout adaptation. No specific rotation angle or sign was repeated consecutively (Figure 1D). Note that the number of trials included in Experiment 2 is the minimum required to ensure that all perturbations are applied to all targets 3 times each, while accounting for the lengthy inter-trial-interval that resulted from aiming to compensate across a larger range of perturbation angles.

Experiment 3 included 10 baseline trials, 20 confidence report practice trials, 150 adaptation trials and 20 washout trials (Figure 1E). The adaptation trials differed from Experiments 1 and 2 in terms of the rotation perturbation applied to the cursor, which was designed to only induce implicit learning. In Experiment 3, trial-by-trial perturbations were pseudo-randomly drawn from a uniform distribution with a range of ±7.5° with a mean value that increases linearly with a step size of 0.33° up to a maximum of 15° (with CW/CCW rotations evenly counterbalanced across participants). By gradually increasing the mean perturbation, the use of explicit aiming strategies was thus constrained (27, 28).

To replicate and generalize the results of Experiment 3, Experiment 4 included the same trial structure as Experiment 3 but with a slightly different perturbation schedule. In Experiment 4, trial-by-trial perturbations were pseudo-randomly drawn from a uniform distribution with a range of ±7.5° centered around a mean of zero degrees, and the mean stayed at zero for the duration of the experiment. The experiment included the same trial structure as Experiment 3 with only one target and no explicit aiming phase. Note that Experiments 3 and 4 had fewer trials compared to Experiments 1 and 2 because they involved reaching to a single target instead of multiple targets.

### Statistical Analysis

Primary dependent variables were confidence judgements and recorded hand angles on every trial. Since participants were instructed to always adjust the confidence bar by at least one unit, trials where the confidence rating remained at the initial 50% mark were removed (Exp. 1: 353 out of 5,184 trials [7.39%]; Exp. 2: 390 out of 4,480 trials [8.71%]). Excluding these trials did not change the identity of the winning model, nor did it significantly change any of the model parameters (all Wilcoxon rank sum tests comparing parameters between exclusion and non-exclusion model fits were insignificant [p=0.60-0.99]). Data was analyzed using Matlab (Mathworks, Inc. Version 2022a). Model fits were computed using Matlab’s *fmincon* function, minimizing the SSE between our confidence models and the confidence report data. Violin plots were generated using the *Violinplot* function in Matlab (29). Data and analysis code can be accessed at https://github.com/ilestz/confidence_analysis.

We validated model parameter optimization through parameter recovery and found we could achieve stable parameter fits throughout. To do this, we fit model-predicted confidence reports using the model that initially generated these predictions and found that model parameters were recovered to 100% accuracy within 2 iterations of fitting. R^2^ values were computed through linear regression of model predictions and data. Reported ΔAIC (30) values reflect differences in summed AIC values between each model and the winning model.

### Computational Modeling

The goal of our study was to examine how motor learning relates to one’s metacognitive judgments of their movement decisions. Participants performed a standard sensorimotor adaptation task while also reporting their confidence in each of their movements (i.e., chosen reach directions; Figure 1). We constructed computational models with the goal of predicting these subjective confidence reports on each trial. All four models we tested characterized confidence reports (Equation 1) as deviations from a maximum confidence ‘offset’ that is proportional to an estimate of previous sensorimotor error(s):

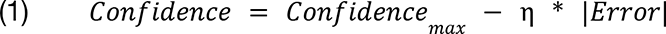

Where η represented an error scaling parameter. Here, “Error” denotes the experienced “target error” (henceforth TE), the absolute magnitude of the angular error of the cursor relative to the target.

The first class of models we tested attempt to predict confidence based simply on the *current state of learning*. Specifically, these models relate confidence reports directly to the most recent error signal, representing a local “one-trial-back” (OTB) update rule. Within this model class, we tested two model variants: In the objective-error one-trial-back model (OTB_obj_), the true (i.e., actually observed) absolute error of the cursor relative to the target was used to compute confidence. Importantly, previous work has shown that the cost of target errors are scaled subjectively via an approximate power-law (31). Thus, we also fit a subjective-error one-trial-back model (OTB_subj_), which scaled all target errors by an exponential free parameter, γ:

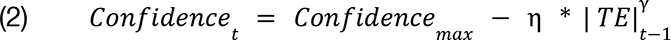

An exponent γ > 1 suggests that large target errors are perceived as relatively more salient (costly), and thus drive sharper decreases in confidence versus small errors in a manner that is not linearly proportional to the veridical error. In contrast, 0 < γ < 1 suggests that large errors are discounted relative to their veridical magnitude and thus drive weaker decreases in confidence than would be predicted by the objective error model. Finally, γ = 1 reduces to the objective error case, where errors are not subjectively scaled and confidence is linearly proportional to the veridical target error magnitude.

The second class of models involved retaining a sort of error memory via estimating a running average of target errors across trials. This estimate is then used to generate predicted confidence reports. We designed these so-called “error-state-space” models to function as simple linear dynamical systems that update an estimate of the current error “state” on every trial through a canonical delta rule:

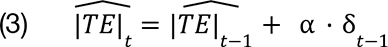

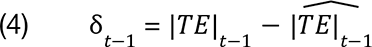

In effect, this learning rule constructs an average of the error state across trials, and echoes the learning rule employed in instrumental learning contexts for learning the predictive value of a given stimulus (32) or state-space models used to model motor adaptation itself (33). The learning rate α reflects the degree to which errors on previous trials are incorporated into the estimate, with high α values (i.e., close to 1) reflecting a high degree of forgetting and low α values (i.e., close to 0) reflecting a more historical memory of error across trials. Whenever the observed target error on the previous trial was greater than (less than) the estimated target error, the estimated target error would increase (decrease) by an amount proportional to this “metacognitive” prediction error.

Within this error-state-space (ESS) model class, there were again two distinct variants: The objective-error state-space model (ESS_obj_) computed the estimated error using the true veridical target errors, while the subjective-error state-space model (ESS_subj_) computed the estimated error using the subjectively scaled error with exponent γ:

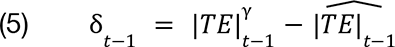

Where the estimated error tracks a history of subjective errors, instead of objective errors. Nonetheless, both history (ESS) models used the same equation to generate predicted confidence reports (Equation 1), now using an evolving estimate of error state:

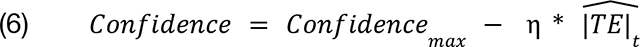

Altogether, our 4 models share 2 free parameters – the maximum confidence offset and the error sensitivity scaling parameter η. Moreover, both models with nonlinear subjective error cost functions share the γ parameter. Finally, both models with error state-space tracking share an additional learning rate free parameter (α) relative to the one-trial-back models.

### Exploratory Regression Analysis

We performed an additional set of regression analyses to ask if confidence computations have downstream effects on learning. To that end, we fit general linear models (GLMs) to changes in hand angle (all experiments) and changes in explicit aim reports (Experiments 1 and 2) from trial to trial. The regressions were designed follows:

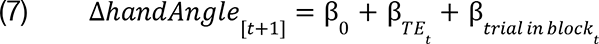

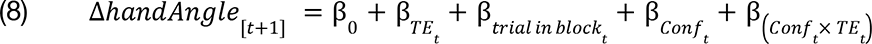

Equation 7 denotes the first GLM that acted as a baseline model to capture changes in movement kinematics from trial *t* to trial *t+1* using the TE on trial *t* and the current trial in the rotation block (or mini-block for Experiment 2) as predictors. Equation 8 denotes an expanded GLM in which confidence ratings (*Conf*) on trial *t* were added as an additional predictor, as well as the interaction between confidence rating and TE. For Experiments 1 and 2, additional variants of the above models were run on changes in explicit aim reports rather than reach directions. Signs of kinematic (or aim) changes were computed relative to the sign of the previous error, such that positive deltas reflected the correct adaptation direction. In Experiments 1 and 2, where participants occasionally skipped the rating phase and left the rating at 50, we removed those trials from the regression analysis. Additionally, changes in hand angle (or explicit Aim for Experiments 1 and 2) that were greater than 3 standard deviations from their mean or were > 20° in the non-adaptive direction (i.e., likely “sign errors”; (34) were removed. GLMs were fit to each subject using the *glmfit* function in MATLAB. Resulting model fits were compared using AIC to account for the increased model complexity in Equation 8.

## RESULTS

### Experiment 1

We sought to explore how participants form subjective judgements of confidence during a motor learning task (Figure 1). Prior to performing a center out reaching motion, participants reported their intended reach direction and rated their confidence that their movement would be successful on a continuous scale (Figure 1B). After a brief baseline phase with veridical cursor feedback, a sensorimotor perturbation of 30° was applied (Figure 1C), which subjects rapidly learned to compensate for. Reach directions compensated well for the applied rotation, with a mean cursor error over the last 50 rotation trials of 0.51° (SD: 3.4°) (Fig. 2A). On average, reach directions compensated for 90% of the perturbation after ∼8 trials and 99% (29.83° [SD: 3.08°]) averaged over the last 20 trials of the adaptation phase. The retention ratio (first 5-trial average of washout divided by the last 5-trial average of adaptation) was 0.38 (SD: 0.14) (t(17) = 11.26, p = 2.6×10-9).

**Figure 2.**
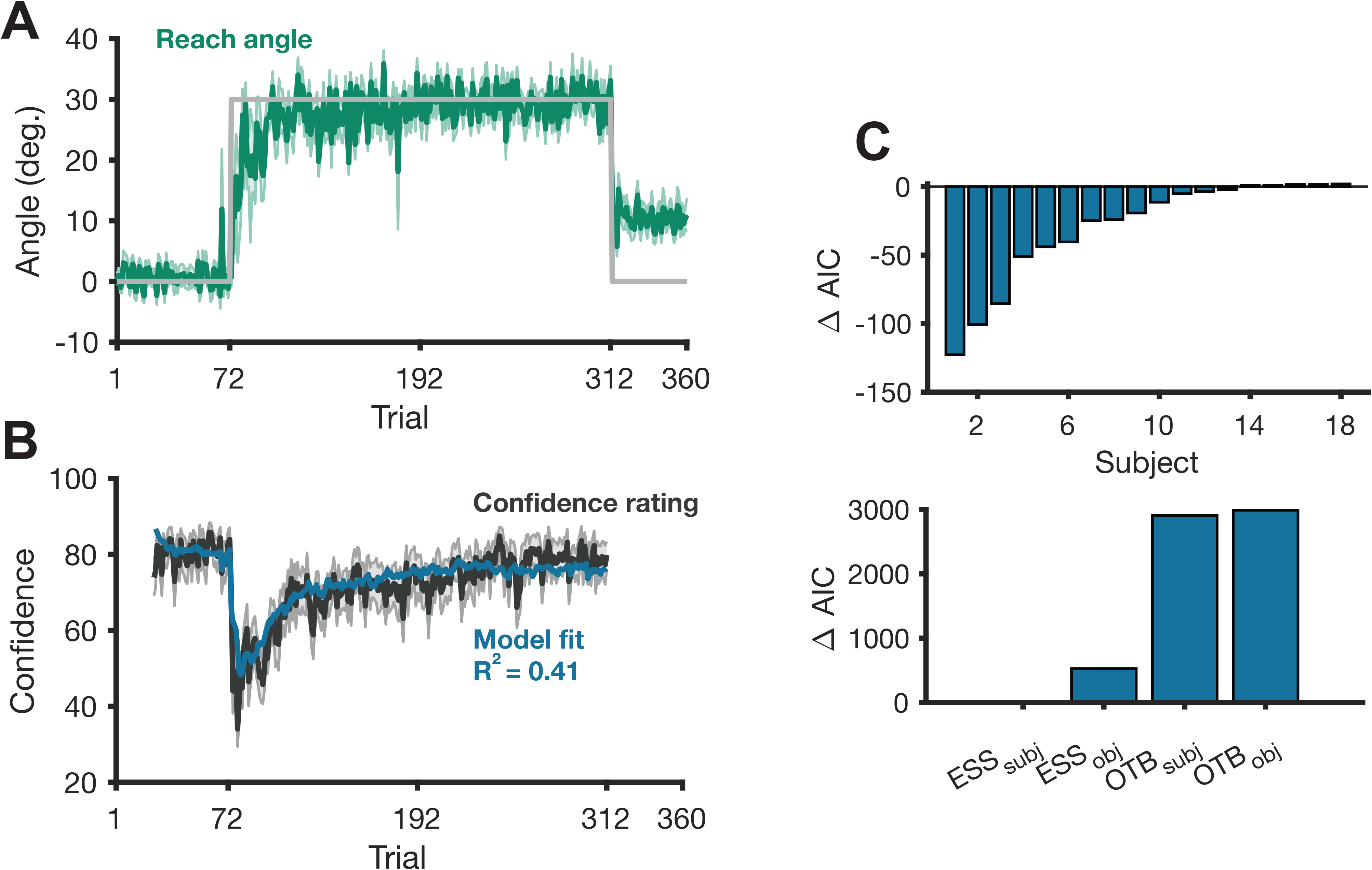
Learning curves and model fitting for Experiment 1 (N=18). (A) Learning curve. Participants adapted their reach angle in response to the 30° perturbation, and showed significant aftereffects in a washout phase. (B) Mean confidence reports (black) and winning model fit (blue). (C) *Top*: AIC differences between the best and second best model are shown for each subject (negative values represent better fits for the winning model, ESS_subj_). *Bottom*: Summed AIC values relative to the summed AIC value for the winning model. All error shading = 1 S.E.M.

During the unperturbed baseline phase of the experiment, confidence reports remained relatively stable, as expected (79.25 [SD:14.67] averaged over all 48 baseline trials). When the perturbation was applied, confidence reports sharply decreased to 45.31 (SD: 21.43 average over the first 5 trials of the adaptation phase). Unsurprisingly, all of our models of confidence were able to account for this decrease as they were all sensitive to error. Following the initial decrease in confidence, all participants gradually restored confidence to near baseline levels as their reaching errors decreased by the last twenty trials of adaptation (Baseline confidence: 79 [SD: 15]; Confidence in the last 20 trials: 78 [SD: 16]; t(17)=0.17; p=0.87) [Fig. 2B], which was also captured by all models. These expected observations provide initial support for the general form of Equation 1, where confidence is proportional to error.

To get a better picture of the dynamics of subjects’ metacognitive judgments, we turned to model comparisons. To reiterate the models tested (see *Methods* for more detailed descriptions): one class of models, the “one-trial-back” models (Equation 2), predicted confidence reports on a given trial based on the current state of learning. The objective-error one-trial-back (OTB_obj_) model predicted confidence based on veridical absolute cursor errors relative to the target, and the subjective-error one-trial-back (OTB_subj_) model predicted confidence based on errors which were scaled by a power-law. The second class of models, the “error-state-space” models, kept track of an estimate of the average error on recent trials and used this average error to compute predicted confidence reports. Again, this class of models either kept track of objective errors (ESS_obj_ model) or subjectively-scaled errors (ESS_subj_ model).

Of the four confidence models we developed, the ESS_subj_ best explained the variance in confidence reports (R^2^=0.41±0.23 [mean±SD]). At the individual level, 14 out of 18 total subjects were better fit by the winning model versus the second best model (Fig. 2C). Moreover, the ESS class of models robustly outperformed the OTB class (Fig. 2C; Table 1). This suggests that metacognitive judgments during sensorimotor learning incorporate a continually-updated history of recent errors, rather than simply acting as a “read-out” of the current state of learning. Furthermore, confidence was better explained by using a subjective error term rather than an objective one: The ESS_subj_ model better fit the confidence versus the ESS_obj_ model (Table 1). All model comparisons were robust, with AIC differences relative to the best-fitting model all exceeding 500.

**Table 1.** Model Fit R^2^ and ΔAIC Values in Experiment 1. ΔAIC are summed AIC values relative to the winning model. Abbreviations: AIC, Akaike Information Criterion; SD, standard deviation; ESS, error-state-space models; OTB, one-trial-back models.

| Model | $R^2$ (SD) | $\Delta AIC$ |
| --- | --- | --- |
| ESS<sub>subj</sub> | 0.4066 (0.2321) | --- |
| ESS<sub>obj</sub> | 0.3587 (0.2272) | 525.5 |
| OTB<sub>subj</sub> | 0.1417 (0.0667) | 2943 |
| OTB<sub>obj</sub> | 0.1208 (0.0540) | 3038 |

While the results of Experiment 1 clearly favored the ESS_subj_ model, some limitations remained. First, because the rotation was of a single value (30°) and was fixed throughout the adaptation phase, the task was relatively easy. Thus, it was important to test if our modeling results generalized to a more complex learning environment, one where both errors and confidence reports would be more variable. Moreover, because of the nature of the task in Experiment 1, both learning curves and confidence reports monotonically increased together; a more variable environment would thus also help us rule out potential coincidental similarities in autocorrelation structure between our winning model and subjects’ learning curves as the key factor. To that end, in Experiment 2 we implemented a pseudo-randomly varying perturbation schedule. This allowed us to both control for the aforementioned limitations, while also testing a novel question we formed *a priori* – are the dynamics of metacognitive confidence judgments during sensorimotor learning affected by environmental uncertainty? Following the logic of control models like the Kalman filter, we expected a shallower history of errors to be integrated into the confidence computations, exemplified by the alpha parameter of our models (Eq 3).

### Experiment 2

Experiment 2 involved perturbations that fluctuated every few trials (i.e., the perturbation changed size and direction every 4 or 8 trials, see Figure 3A and *Methods*). This allowed us to perform a more strict test of our modeling approach, and to examine if and how environmental uncertainty affected subjective confidence reports. Specifically, we predicted that the ESS_subj_ model would best account for subjective confidence ratings in this context, replicating Experiment 1. Moreover, we also hypothesized that while the fundamental process of confidence ratings would remain the same (i.e., the ESS_subj_ would again best explain behavior), the learning rate parameter of that model would increase in response to the increase in environmental uncertainty such that it would incorporate a more recency-biased history of errors (17).

**Figure 3.**
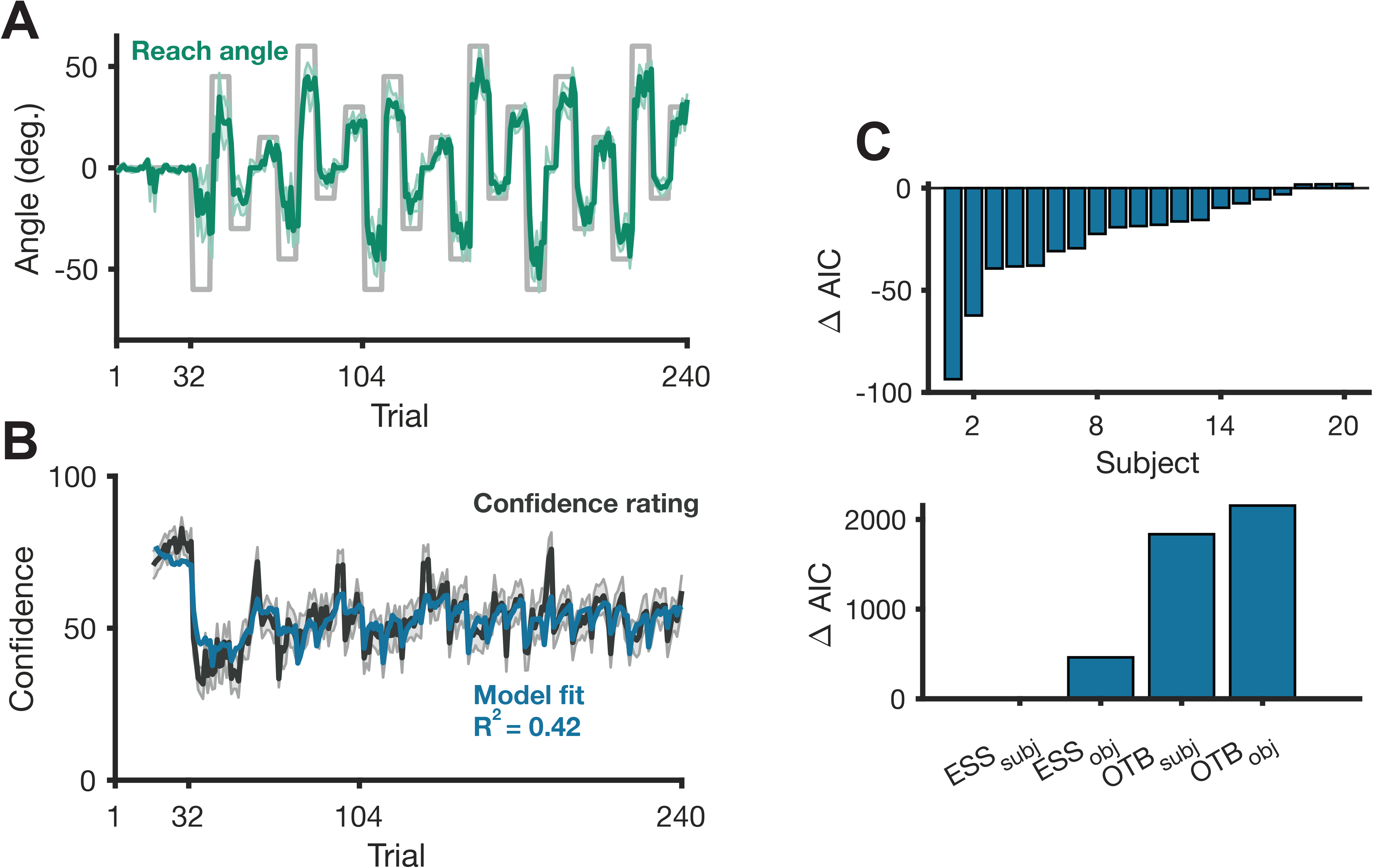
Learning curves and model fitting for Experiment 2 (N=20). (A) Learning curve. Participants adapted their reaches (green) to account for the volatile perturbation schedule (gray). (B) Mean confidence reports (black) and winning model fit (blue; ESS_Subj_). (C) *Top*: AIC differences between the best and second best model are shown for each subject (negative values represent better fits for the winning model, ESS_Subj_). *Bottom*: Summed AIC values relative to the summed AIC value for the winning model. All error shading = 1 S.E.M.

Despite the more volatile nature of the perturbation schedule, participants were still able to alter their reach directions to account for the rotations (Fig. 3A). Excluding the transition trials where the rotation abruptly changes, subjects’ average cursor error in the last two perturbation blocks was only 6.3° (SD: 6.5°). A GLM using block number, trial position within a block, and their interaction to predict percent compensation for each subject found that trial position within a block and the interaction term both significantly predicted percent compensation. Specifically, later trials within a block elicited greater compensation for the perturbation (β=0.08 [SD: 0.06], t(19)=6.28, p=4.9×10^-6^), indicating learning within a block. Additionally, the interaction term revealed that participants exhibited faster learning in later blocks (β=0.002 [SD:0.003], t(19)=3.03, p=0.007), supporting the presence of “meta-learning.”

As in Experiment 1, confidence remained relatively stable during the unperturbed baseline but sharply decreased after the onset of the first perturbation (Fig. 3B). Confidence also tended to sharply decrease at the start of each new perturbation. Throughout the experiment, some 4-trial zero-rotation blocks were introduced, and these blocks tended to coincide with high confidence reports (Fig. 3B).

Once again, the ESS_subj_ model best predicted confidence reports in this experiment (R^2^=0.42±0.20 [mean±SD]), and the one-trial-back models were unable to account for the large fluctuations in confidence reports and performed significantly worse (Table 2). At the individual level, 18 out of 20 total subjects were better fit by the winning model versus the second best model (Fig. 3C). Both history (ESS) models again tracked confidence reports more accurately than the other class of models (OTB). Thus, our model comparison results closely replicated those of Experiment 1 (and again were robust; lowest AIC difference: 498). This further suggests that metacognitive judgements of sensorimotor learning incorporate a gradually changing history of (subjectively-scaled) errors. We do note that none of the four models were able to fully capture the unusually high confidence ratings seen during the zero-rotation blocks (see *Discussion*). Finally, the alpha parameter significantly increased when comparing Experiment 2 to Experiment 1, inline with our hypothesis and normative predictions concerning the effect of volatility on learning rates (Exp. 2: 0.46 [SD:0.23]; Exp. 1: 1.91 [SD:0.93]; Wilcoxon rank sum test; Z=3.29, p=5.3×10^-7) (17). Additional analysis of model parameters is given in the section *Comparing model parameters across experiments*.

**Table 2.** Model Fit R^2^ and ΔAIC Values in Experiment 2. ΔAIC are summed AIC values relative to the winning model. Abbreviations: AIC, Akaike Information Criterion; SD, standard deviation; ESS, error-state-space models; OTB, one-trial-back models.

| Model | $R^2$ (SD) | $\Delta AIC$ |
| --- | --- | --- |
| ESS<sub>subj</sub> | 0.4207 (0.1974) | --- |
| ESS<sub>obj</sub> | 0.3539 (0.1848) | 498.1 |
| OTB<sub>subj</sub> | 0.2373 (0.1009) | 1984 |
| OTB<sub>obj</sub> | 0.1741 (0.0830) | 2316 |

### Experiment 3

The inclusion of an abrupt, large perturbation and the requirement to aim promoted the use of explicit strategies in Experiment 1. The inclusion of multiple perturbations with different sign and magnitude across repeating blocks in Experiment 2 constrains implicit learning and also promotes the use of explicit strategies. As such, it is difficult for us to disentangle whether confidence tracks improvements in the precision of explicit aiming strategies, unmeasured proprioceptive errors between the desired aim and the kinesthetic feedback from the arm, or simple the observed visuomotor target errors, as we assumed in our modeling. Thus, we performed two additional Experiments designed to target implicit learning alone. If confidence mainly tracks a generic target error signal, our model should continue to perform well in this context; however, if confidence mainly responds to changes in explicit motor learning, it should not. Experiment 3 sought to eliminate explicit aiming strategies by imposing a noisy rotation whose mean gradually increased over 45 trials to 15°, but which had uniform noise of ±7.5°around the mean. This perturbation schedule should generally preclude any explicit aiming strategies as participants would not be aware of the gradually increasing mean perturbation (31–33) nor can predict the noise. We hypothesized that the ESS_subj_ model would still be able to explain fluctuations in confidence, supporting the assumption that confidence tracks observed target errors.

Despite the randomness of the perturbation and the lack of explicit aiming strategy, participants implicitly adapted well to the mean of the perturbation, reaching an average of 85.62% (SD: 19.46%) compensation by the last 5 trials of the adaptation phase (Fig. 4A). Significant after effects were observed (retention ratio: 0.61 [SD:0.80]; t(23)=3.66; p=0.001), consistent with the implicit nature of adaptation to this perturbation. While participants’ adaptation largely tracked the mean of the perturbation, their subjective confidence reports slightly decreased over the course of the experiment (Fig. 4B, *upper*; confidence in the first 20 rating trials: 63.7 [SD:17.3]; confidence in the last 20 rating trials: 51.0 [SD: 22.9]; t(23)=2.34; p=0.03). This decrease in confidence appeared to reflect increased cursor errors (cursor error in the first 20 trials: 2.80° [SD: 2.36°]; cursor error in the last 20 trials: 4.72° [SD: 2.15°]; t(23)=7.67, p=8.7×10^-8^).

**Figure 4.**
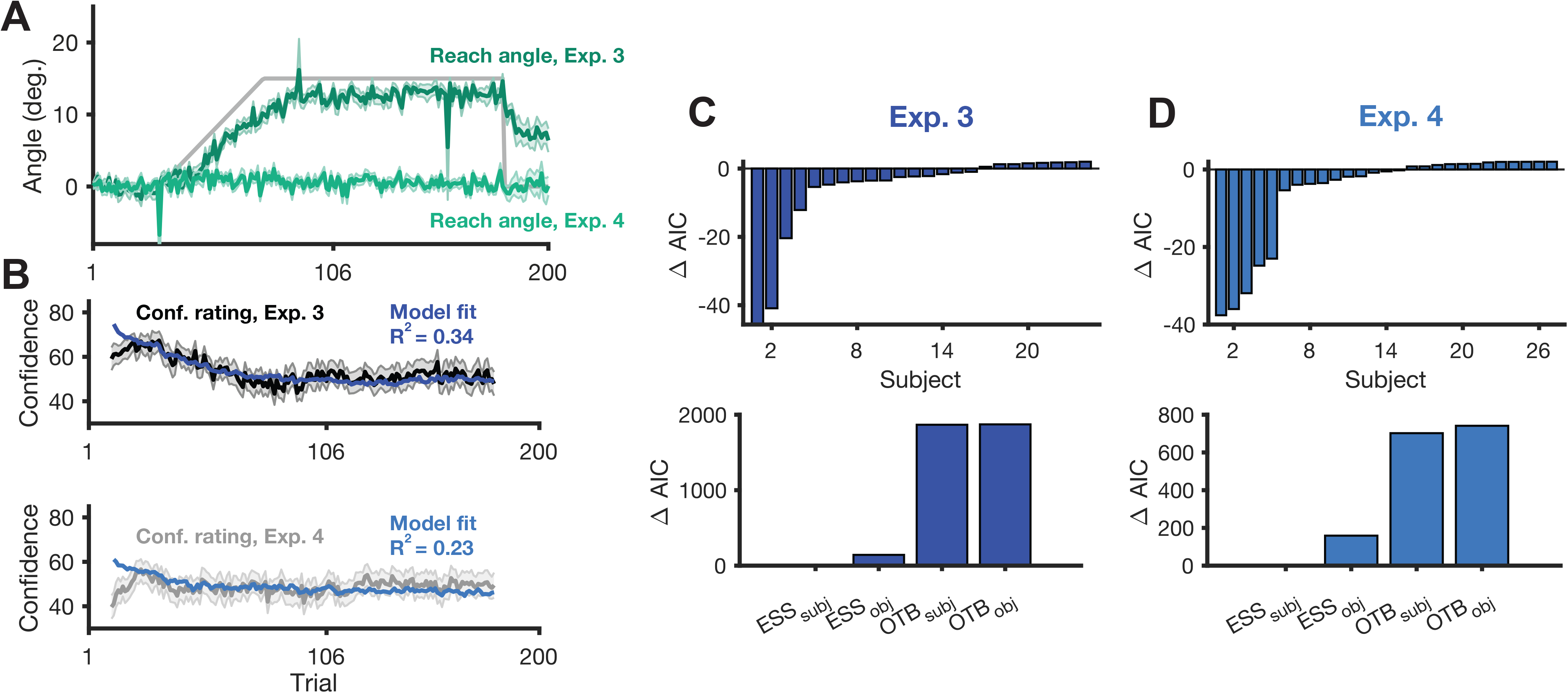
Learning curves and model fitting for Experiments 3 (N=25) and 4 (N=30). (A) Learning curve. Participants adapted their reaches (green) to account for the volatile perturbation schedule (gray). (B) Mean confidence reports (black and gray) and winning model fit (dark and light blue; ESS_Subj_). (C-D) *Top*: AIC differences between best and second best model are shown for each subject (negative values represent better fits for the winning model, ESS_Subj_). *Bottom*: Summed AIC values relative to the summed AIC value for the winning model. All error shading = 1 S.E.M.

As hypothesized, the ESS_subj_ model best fit subjects’ confidence ratings, achieving the lowest summed AIC and a mean R^2^ of 0.33 (SD: 0.34) (Fig. 4C; Table 3). The ESS_subj_ model best fit 11 out of 24 subjects. Of the 14 remaining subjects, 7 were best fit by the ESS_obj_ model, suggesting that the exponent parameter was largely unnecessary for these subjects. Overall 18 out of 24 subjects were better fit by history (ESS) models, thus replicating the finding in Experiments 1 and 2 that the ESS models best fit the data over the one-trial-back (OTB) models and that confidence tracks a history of performance errors.

**Table 3.** Model Fit R^2^ and ΔAIC Values in Experiment 3. ΔAIC are summed AIC values relative to the winning model. Abbreviations: AIC, Akaike Information Criterion; SD, standard deviation; ESS, error-state-space models; OTB, one-trial-back models.

| Model | $R^2$ (SD) | $\Delta AIC$ |
| --- | --- | --- |
| ESS<sub>subj</sub> | 0.3327 (0.3405) | --- |
| ESS<sub>obj</sub> | 0.3047 (0.3385) | 240 |
| OTB<sub>subj</sub> | 0.0704 (0.0997) | 2271 |
| OTB<sub>obj</sub> | 0.0586 (0.0825) | 2284 |

### Experiment 4

Building on the findings of Experiment 3, Experiment 4 aimed to further disentangle the contributions of explicit strategies and implicit learning processes to confidence judgments. Here, by introducing a zero-mean perturbation, we eliminated any correlation of confidence with steady improvements in motor performance; however, by applying a noisy distribution of perturbations we again precluded the use of explicit aiming strategies, providing us a context in which to further validate our model predictions.

Given that the perturbation was randomly sampled from a uniform distribution ranging from -7.5° to +7.5°, participants could not predict the rotation on any given trial. However, the resultant cursor errors would produce a small degree of single trial implicit learning. On average, participants adjusted their hand angle by 1.58° (SD: 1.47°) in the opposite direction of the cursor error on the subsequent trial. This single trial learning was significantly different from zero (t(26)=5.56, 7.8×10^-6^), indicating some degree of implicit adaptation. Confidence ratings remained largely stable throughout the experiment (Fig. 4B, *lower*; confidence in the first 20 perturbation trials: 50.2 [SD: 22.8]; confidence in the last 20 perturbation trials: 50.0 [SD: 25.2]; t(26)=0.05; p=0.96), as did average performance (Fig. 4A; cursor error magnitude in the first 20 perturbation trials: 4.04° [SD: 1.33°]; cursor error magnitude in the last 20 perturbation trials: 4.24° [SD: 0.98°]; t(26)=-0.83, p=0.41).

As in Experiment 3, the ESS_subj_ model again gave the best fit to subjects’ confidence ratings, achieving the lowest summed AIC and a mean R^2^ of 0.24 (SD: 0.21) (Fig. 4D; Table 4). The ESS_subj_ model best fit 13 out of 27 participants. Of the remaining 14, 8 participants were best fit by the ESS_obj_ model, as the exponent parameter for these participants was close to 1 (and thus less important). Overall, 77.8% of participants (21/27) were better fit by an error history (ESS) model, rather than a one-trial-back (OTB) model. Taken together, the results of Experiments 3 and 4 suggest that confidence during motor learning reflects a form of performance monitoring that tracks performance error history regardless of whether learning is driven by explicit changes in movement strategy or implicit adaptation.

**Table 4.** Model Fit R^2^ and ΔAIC Values in Experiment 4. ΔAIC are summed AIC values relative to the winning model. Abbreviations: AIC, Akaike Information Criterion; SD, standard deviation; ESS, error-state-space models; OTB, one-trial-back models.

| Model | $R^2$ (SD) | $\Delta AIC$ |
| --- | --- | --- |
| ESS<sub>subj</sub> | 0.2316 (0.2055) | --- |
| ESS<sub>obj</sub> | 0.2204 (0.2091) | 126 |
| OTB<sub>subj</sub> | 0.1029 (0.1339) | 764 |
| OTB<sub>obj</sub> | 0.0883 (0.1227) | 794 |

### Comparing model parameters across experiments

While the variance in confidence reports was best explained in all experiments by the subjective-error history (ESS_subj_) model, parameter values in each experiment were not expected to be the same. In fact, given the differences in perturbation schedules across experiments, two key parameter value differences were hypothesized. First, as discussed above, it has been observed in other fields that greater environmental volatility results in higher learning rates (17, 38). Thus, we predicted that Experiment 2 should have a higher learning rate (α) compared to other experiments, as the perturbation abrupted changed several times throughout the task. Although the degree of perturbation noise in Experiments 3 and 4 was also high, we do not predict a large learning rate *per se* as the noise was consistent throughout the task, drawn from a uniform distribution, in contrast to the sudden changes of perturbations applied in Experiment 2.

We also predicted that subjective error computations would be sensitive to the general error distribution induced by the task. Since Experiments 2-4 exposed participants to much larger errors on average than Experiment 1, we would expect smaller exponent γ values, reflecting the hypothesis that participants would discount frequent large errors. Conversely, in Experiment 1, since most errors were small, after the initial onset of the perturbation we would expect a larger exponent γ value, reflecting increased salience of small errors. An inherent trade-off in our models also predicts that smaller exponent γ values induce increased sensitivity parameter η values. Consistent with this trade-off and the above prediction regarding γ values, Experiment 1 would be predicted to have the smallest sensitivity parameter η.

Due to the large number of comparisons conducted while investigating parameter differences across experiments, we controlled Type-1 error rates by adjusting p-values using the false discovery rate procedure (39). Learning rates were significantly higher in Experiment 2 relative to Experiments 1 and 3 (Fig. 5D; Wilcoxon rank sum test; Exp. 1: Z=3.29, p_adj_=0.006; Exp. 3: Z=2.65, p_adj_=0.02), largely in line with our hypothesis. However, there was no significant difference in learning rates between Experiments 2 and 4 (Wilcoxon rank sum test; Z=1.41; p_adj_=0.23).

**Figure 5.**
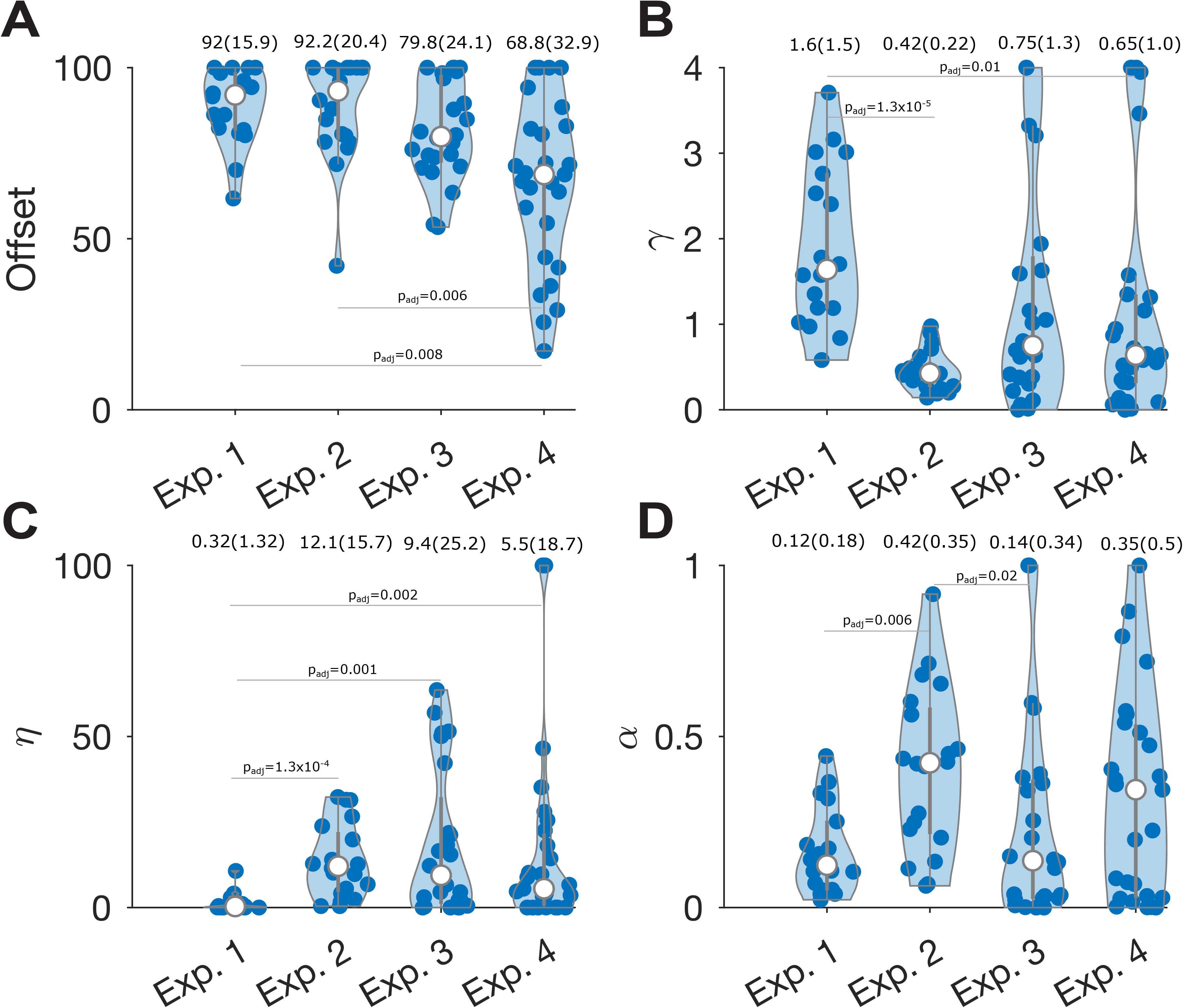
Parameter comparison across Experiments. Green shaded regions reflect the distribution of participant parameter values. White dots indicate median parameter values and gray bars the interquartile range (IQR) between the first and third quartiles. Median and IQR values are included above each plot and Wilcoxon adjusted p-values are provided in-situ. Whiskers extend to 1.5 times the IQR. (A) Maximum confidence offset parameters. (B) Subjective scaling of errors via an exponential free parameter. (C) The sensitivity parameter. (D) Confidence model learning rates.

Our second hypothesized parameter difference across experiments was that Experiment 1 would show a higher exponent γ and consequently a lower sensitivity η relative to other experiments where errors tended to be larger. Experiment 1 had a significantly lower sensitivity η compared to all three other experiments (Fig. 5C; Wilcoxon rank sum test; Exp. 2: Z=-4.40, p_adj_=1.3×10^-4^; Exp. 3: Z=-3.04, p_adj_=0.01; Exp. 4: Z=-2.63, p_adj_=0.02). Experiment 1 also showed a higher exponent γ compared to Experiments 2 and 4 (Fig. 5B; Wilcoxon rank sum test; Exp. 2: Z=5.01, p_adj_=1.3×10^-5^; Exp. 4: Z=2.86, p_adj_=0.01), but the comparison to Experiment 3 did not survive corrections (Wilcoxon rank sum test, Z=2.12, p_adj_=0.07). These findings are mostly consistent with our hypothesis that in Experiments 2-4, where errors tended to be larger, participants discounted larger errors to a greater degree than they did in Experiment 1, where errors tended to be smaller.

While we did not have *a priori* predictions about the maximum confidence offset parameter, analyses revealed that Experiment 4 had a significantly lower offset than Experiments 1 and 2 (Fig. 5A; Wilcoxon rank sum test; Exp. 1: Z=-3.14, p_adj_=0.008; Exp. 2: Z=-3.37, p_adj_=0.006) but the comparison to Experiment 3 did not survive corrections (Exp. 3: Z=-2.27, p_adj_=0.055). The maximum confidence parameter reflects the theoretical maximum confidence a participant would rate if they had no cursor error on any trial. If participants are using the full range of the scale, this parameter should be close to maximal (i.e. 100). However, in Experiment 4, the mean parameter value was 66 (SD: 24). Participants in Experiment 4 largely had very low confidence ratings and rarely rated highly on the scale, likely due to the unpredictable noisy perturbation, so it perhaps is reasonable that their maximum confidence parameter would be lower than the other experiments. Taken together, our between-experiment parameter results (Fig. 5) suggest that subjects adapted the dynamic range and timescale of error integration of their confidence computations in a manner that adapted to the statistics of the environment.

### Does confidence affect motor learning?

Thus far, our findings have examined the effect of errors on participants’ subjective confidence. While it may seem intuitive that confidence ratings incorporate information about recent motor performance, confidence judgements may also modulate changes in motor behavior, rendering confidence ratings as more than just an epiphenomenal “read-out” of task-error. Indeed, confidence in our task reflects a form of subjective (un)certainty in one’s actions, which according to Bayesian logic may influence how someone behaves (40, 41). Does having high or low confidence predict anything about participants’ ability to adapt to errors? In other words, is there a bi-directional relationship between confidence and motor learning?

To address this we explored whether confidence on trial *t* could predict changes in motor behavior on trial *t+1*, above and beyond the changes predicted by error alone. We fit generalized linear models (GLMs) to the kinematic data on each trial (see *Methods* for details). The first GLM acted as a control, only attempting to predict changes in hand angle using TE on trial *t,* and the current trial in the rotation block (or mini-block for Experiment 2) as control predictor. The second GLM added the confidence rating on trial *t* as an additional predictor, as well as the interaction between confidence rating and TE.

Intriguingly, in both Experiments 1 and 2 adding confidence to the regression model significantly improved the model’s ability to predict trial-by-trial changes in motor behavior, both when using changes in explicit aim or raw hand angle as the predicted variable (see *Methods*). In Experiment 1, The confidence model fit to changes in aim outperformed the control (i.e., without-confidence) model by an average of 30.3 AIC points (SD: 29.6; Wilcoxon sign rank test on differences in AIC values across the sample, Z=-3.46, p=5.4×10^-4^). Regression coefficients for the confidence term did not significantly differ from zero (Z=-1.28, p=0.20). Similar results were seen when using raw hand angle changes as the predicted variable (which we note were correlated with aim changes) – the confidence model used to predict changes in hand angle also outperformed the control model, by an average of 21.7 AIC points (SD: 26.4; Wilcoxon sign rank test, Z=-3.20, p=0.001). Regression coefficients for the confidence term again did not differ from zero (Z=0.68, p=0.41). In Experiment 2, confidence did significantly predict changes in aim, improving the model fit by an average of 19.1 AIC points (SD: 19.9; Wilcoxon sign rank test, Z=-3.55, p= 3.89×10^-4^). Unlike in Experiment 1, the regression coefficient for the confidence term was significantly different from zero (Mean:-1.82; SD: 1.86; Z=-3.36, p= 3.9×10^-4^). This suggests that changes in aim following a given error were larger if the error occurred on a trial where the participant’s confidence was low. The results were similar when modeling changes in hand angle, which we note were highly correlated with changes in aim (ΔAIC=-22.0 [SD: 26.8], Wilcoxon sign rank test, Z=-3.47, p=5.2×10^-4^). The regression coefficient for the confidence term used to predict changes in hand angle was again significantly less than zero (Mean:-1.41; SD: 2.32, Z=-2.28, p=0.02). This again suggests that changes in hand angle in response to a given error were larger if the error occurred on a low confidence trial. Speculatively, this may be consistent with a Bayesian account – if you have a highly “certain” prior that you will perform well on a trial, sensory error may be somewhat discounted, potentially leading to a smaller change in response to an observed sensory error.

In Experiments 3 and 4, where explicit aiming strategies were largely absent, confidence no longer significantly improved model fits to changes in motor behavior. In fact, in both experiments, the confidence regression model performed significantly worse than models using error alone due to the AIC penalty for additional parameters (Wilcoxon sign rank test; Exp. 3: Z=2.60, p=0.009; Exp. 4: Z=4.06, p=4.9×10^-5^, both in favor of the control model). Thus, in the absence of an explicit component of motor learning, it appears that subjective confidence (certainty) in one’s movements no longer modulates adaptation to errors. This suggests that implicit motor adaptation may be largely impervious to metacognitive judgments of performance. On the other hand, the regression analysis in Experiments 1 and 2 indicate that confidence may modulate motor learning through changes in volitional movement strategies.

## DISCUSSION

Here we examined the relationship between subjective confidence judgments and motor errors in the context of sensorimotor adaptation. We investigated this relationship via four sensorimotor learning experiments that differed with respect to environmental volatility and the perturbation applied, each eliciting varying degrees of explicit or implicit motor learning. We constructed computational models with the goal of predicting subjective confidence reports on each trial based on people’s error histories. We specified a set of models where trial-by-trial subjective confidence tracked only the current learning state (i.e., the most recent performance error), and another set of models where confidence judgments are approximated by a simple linear dynamic system that tracks a recency-weighted history of errors made during learning.

In all four experiments, the ESS_subj_ model, an error history model that also scales performance errors using a subjective cost function, best accounted for subjects’ confidence data. In Experiment 1, this model was best able to account for the confidence data in the context of a fixed, abrupt perturbation schedule. In Experiment 2, subjects learned in a volatile context, and the ESS_subj_ model was again best able to account for the large fluctuations in confidence reports we observed. In Experiments 3 and 4, where the perturbation was small and the perturbation schedule was largely random and thus elicited primarily implicit adaptation, the error history models once again best fit the confidence data. Taken together, these findings demonstrate that confidence computations during sensorimotor adaptation approximate a running average of recent (subjectively-scaled) performance errors. To wit, these findings suggest that when people make metacognitive judgments of their own state of sensorimotor learning, they incorporate a recent history of performance errors rather than just taking a snapshot of their current performance state.

Notably, comparing model parameters between experiments provided additional key insights into the dynamics of metacognitive judgements of performance during sensorimotor adaptation, showing that confidence computations adapt to the statistics of the learning environment. The subjective costs of errors, reflected in the exponent term of our model, varied in accordance with the range of errors experienced by participants in all four experiments. Importantly, the subjective error cost function across experiments agrees with prior observations that small errors are exaggerated and large errors are discounted (31). Moreover, environment volatility affected confidence computations. In Experiment 2, the perturbation abruptly changed across short mini-blocks, resulting in a highly volatile context and a large contribution from explicit learning processes. In line with work in other fields (13), this prompted a higher learning rate in the confidence model fit to Experiment 2. Future studies that parametrically alter aspects of the learning environment (e.g., consistency and variability, [39]) and measure their effects on metacognitive judgements could be useful for developing more detailed models of confidence computations during motor learning.

Despite all four experiments eliciting varying degrees of explicit and implicit motor learning, the winning model was consistent across these contexts. In Experiments 1 and 2, learning was dominated by explicit strategies to counter the rotation. On the other hand, in Experiments 3 and 4, explicit strategies were absent, and implicit learning dominated. Nonetheless, confidence judgments tracked the recent history of task errors and were generally sensitive to subjective versus objective error scalings. This suggests to us that confidence does not merely “read-out” the progress of explicit or implicit processes, but rather reflects the active monitoring of a history of recently observed task performance errors. Task errors emerge as an interface between implicit and explicit processes, indicating the adequacy of the combined effort of these processes for achieving performance goals (43). This error signal, we propose, is most informative for our subjective formation of confidence in our motor performance.

What are the psychological mechanisms that track the error-state used for metacognitive judgments? Although our models are straightforward and principally descriptive, they do constrain the time-scale at which error-signals are integrated, hinting at a role for short term memory in the process of subjective confidence formation. We speculate that working memory is likely important in the formation of confidence judgments during motor learning (32, 33). That is, participants may track the quality of their performance by storing recent outcomes in working memory and integrating them into an estimate of the “state” of their performance. If this is correct, one prediction is that disrupting working memory may alter the relationship between confidence and recent errors. Future studies, perhaps using dual tasks that tax working memory, could test this prediction.

In addition to being able to predict confidence judgments given the history of performance, an exploratory analysis of our data provided some evidence that subjective confidence can modulate changes in motor behavior. Namely, participants’ degree of confidence in their actions immediately preceding an error can predict the degree to which they adjust their actions in response to those errors. We note that we only observed this relationship in Experiments 1 and 2, where explicit learning dominated, but not in Experiments 3 and 4, where learning was largely implicit. Moreover, in Experiment 2 we observed a consistent relationship between confidence and learning form error, where lower confidence appeared to precede greater sensitivity to errors. It is possible that this echoes a Bayesian integration process, where certainty in one’s actions going into a trial reflects a “prior” on the expected error, sensory error feedback represents an observation of new data, and the stronger the prior (i.e., the higher the confidence/lower the uncertainty) the less adaptation to that error. While we only observed this intriguing effect in Experiment 2, future studies that control sensory uncertainty could help us more directly test this idea.

We note several limitations in our study. First, motor perturbation tasks should not be conflated with true skill learning (24); measuring confidence judgments in more complex motor skill learning tasks will be essential for asking if our models generalize. Second, many models of confidence take a Bayesian approach (16), explicitly modeling sensory uncertainty as a key component of confidence. We took a more descriptive approach here by focusing on the overall dynamics of confidence judgments during visuomotor learning, and by focusing on confidence in one’s performance, not sensory perception. Future studies could also incorporate uncertainty in other forms (e.g., sensory feedback, increased motor noise, etc.) to further develop our models in a more normative framework. Third, participants in Experiment 2 exhibited higher confidence during the zero-rotation blocks, where feedback was similar to the baseline phase of the experiment and was completely unperturbed. One possible explanation for this behavior is that participants may have been able to compare the performance of the cursor to their intended movement, signaling a change in context reminiscent of the veridical baseline experience in earlier trials. This would explain why confidence on the zero-rotation blocks appears to return to baseline levels, as participants already have experience with zero-rotation trials. Importantly, an error signal based on the discrepancy between the aimed movement and cursor is insufficient to explain the variation in confidence observed in, for example, experiment 3, where participants adapt implicitly. Instead, task errors (the observed error of the cursor relative to the center of the target) are a better predictor of confidence across all of our experiments. Thus, confidence in this context quickly returned to baseline levels on these trials. Our current model cannot account for this potential long-term memory effect which may involve familiarity with previous baseline phases, the detection of large perturbations, and contextual shifts.

Moreover, it is likely the case that confidence is related to “agency” (44–46). How is confidence influenced by errors that are perceived to be a consequence of one’s own motor variability versus the environment? This could be another important topic of further study. Finally, our model was less able to capture variance in Experiments 3 and 4 versus Experiment 1 and 2.; in the former, the perturbation schedule was random and involved smaller errors. This may again relate to judgments about agency and the source of errors, though at present we can only speculate as such.

The prospect that higher-level metacognitive judgements accurately track lower-level sensorimotor variables (e.g., visual error magnitudes) compels the search for underlying neural correlates. In the context of sensorimotor learning, confidence can be defined as a higher-order variable that corresponds to the uncertainty that underpins the learning process (18, 19). Multiple neural regions capable of representing sensory uncertainty have been proposed, including the orbitofrontal cortex (OFC) (47, 48), midbrain (49), anterior cingulate cortex (ACC) (50), insula (51, 52), and prefrontal cortex (PFC) (53, 54). In terms of representations of confidence, activity in the rostrolateral and dorsolateral PFC (rlPFC/DLPFC) is purported to be central to the processing of explicit confidence judgements in decision making (55–58). It may be that some of these regions, in addition to areas involved in working memory, could show functional correlations to the variables we have modeled here. That is, our model (or similar ones) could we used in attempts to track or disrupt neural correlates of metacognitive variables during motor learning – such as the estimated error state (Equation 3) or metacognitive prediction errors (Equation 4) – using techniques like fMRI and TMS.

Understanding the potential role of metacognition in tracking motor learning progress, and perhaps exerting an influence on the learning process itself, has several potential practical implications. In clinical contexts, understanding the factors that promote increased confidence in a patient’s motor ability may help predict their persistence in therapy as well as help speed up their performance improvements through treatment. Moreover, there is a rather large body of work on metacognition in childhood education (59–61); if, say, a child is struggling with a motor skill (e.g., handwriting), understanding how their recent progress influences their metacognitive conceptions can be useful knowledge. The same principle can be applied to inform how metacognition plays a role in professional sports, and perhaps inform future coaching approaches.

In conclusion, here we show that a simple Markovian learning rule was able to capture people’s confidence ratings as they performed skilled reaches and adapted those movements in response to errors. Our model showed that people’s metacognitive judgment of their motor performance appeared to incorporate a recency-weighted history of subjectively-scaled sensorimotor errors. This model was robust to different learning contexts that elicited varying degrees of implicit or explicit learning. The specific statistics of the learning environment, including volatility of the perturbation and the average magnitude of errors, lawfully altered how errors were integrated into metacognitive judgments of confidence. Further, we demonstrate that metacognitive confidence computations may exert an influence on explicit motor compensation strategies in response to errors. Our findings provide a foundation for future studies to investigate sensorimotor confidence during more real-world learning tasks, and to localize its correlates in the brain.

## Acknowledgements

We thank Ryan Morehead, Jordan Taylor, Huw Jarvis, and the ACT lab for helpful discussions.

C.L.H was funded by Yale University’s Seesel Postdoctoral Fellowship.

## Captions

**Table T1.** *Note.* T-tests are compared to target error model R^2^, unless stated otherwise. Abbreviations: df, degrees of freedom, SD, standard deviation.

